# Insights into dynamic sound localisation: A direction-dependent comparison between human listeners and a Bayesian model

**DOI:** 10.1101/2024.04.26.591250

**Authors:** Glen McLachlan, Piotr Majdak, Jonas Reijniers, Michael Mihocic, Herbert Peremans

## Abstract

Self-motion is an essential but often overlooked component of sound localisation. While the directional information of a source is implicitly contained in head-centred acoustic cues, that acoustic input needs to be continuously combined with sensorimotor information about the head orientation in order to decode these cues to a world-centred frame of reference. On top of that, the use of head movement significantly reduces ambiguities in the directional information provided by the incoming sound. In this work, we evaluate a Bayesian model that predicts dynamic sound localisation, by comparing its predictions to human performance measured in a behavioural sound-localisation experiment. Model parameters were set a-priori, based on results from various psychoacoustic and sensorimotor studies, i.e., without any post-hoc parameter fitting to behavioral results. In a spatial analysis, we evaluated the model’s capability to predict spatial localisation responses. Further, we investigated specific effects of the stimulus duration, the spatial prior and sizes of various model uncertainties on the predictions. The spatial analysis revealed general agreement between the predictions and the actual behaviour. The altering of the model uncertainties and stimulus duration revealed a number of interesting effects providing new insights on modelling the human integration of acoustic and sensorimotor information in a localisation task.

**Author summary:** In everyday life, sound localisation requires both interaural and monaural acoustic information. In addition to this, sensorimotor information about the position of the head is required to create a stable and accurate representation of our acoustic environment. Bayesian inference is an effective mathematical framework to model how humans combine information from different sources and form beliefs about the world. Here, we compare the predictions from a Bayesian model for dynamic sound localisation with data from a localisation experiment. We show that we can derive the model parameter values from previous psychoacoustic and sensorimotor experiments and that the model without any post-hoc fitting, can predict general dynamic localisation performance. Finally, the discrepancies between the modelled data and behavioural data are analysed by testing the effects of adjusting the model parameters.

## Introduction

The acoustic cues acquired through head rotation are of crucial importance to spatial hearing. Not only do they improve sound externalisation [1], they provide dynamic acoustic cues which contribute to sound localisation. More specifically, they reduce ambiguities between the front and back [2, 3] and, under certain conditions, improve the elevation estimation of a source significantly [4, 5]. Dynamic acoustic cues become especially important when spectral cues are difficult to process, like in reverberant [6] or virtual [7] environments.

The movements that benefit sound localisation are not always voluntary. On the contrary, for both sensorimotor [8] and behavioural [9, 10] reasons, the human head is rarely completely still. Even when the only task given to subjects is to remain still, they continue to move by a small but measurable amount [11]. Despite this constant motion, we perceive the auditory world to be relatively stable. This suggests that there exists a mechanism that utilises positional information about the head to compensate for self-motion and converts the head-centred auditory cues into a stable, world-centred frame of reference [12]. This notion is further supported by the fact that moving sound sources do not provide the same benefits to localisation as the head movements that would theoretically cause similar acoustic cues [13].

Due to their complicated methodology, studies on sound localisation that involve head movement are scarce. It follows that the available work is also less extensive than that on static sound localisation. For example, full direction-dependent analyses of two-dimensional sound localisation have been conducted in the past for static listening [14, 15]. A similar in-depth spatial analysis is not yet available for sound localisation that involves head movement. Most studies restrict the presented source positions to the horizontal or median plane [3, 5]. Studies that do allow stimuli from more directions simplify their analyses by collapsing performance metrics together over several directions [4, 13].

If we are to include these unavoidable dynamic effects in future studies of sound localisation, it is apparent that there is a need for a tool to better investigate or predict the effects of head movements in an easily reproducible manner. To this end, we previously proposed a Bayesian framework to model dynamic sound localisation through self-motion by integrating acoustic and sensorimotor information over time [16]. This model provides a bottom-up approach to sound localisation, i.e., it lets one change the cues that are extracted from incoming sound and the way that they are utilised (e.g. by choosing the spatial prior or decision rule), after which the effects on localisation performance can be tested.

In this paper, we quantitatively validate the proposed Bayesian model for dynamic sound localisation, by comparing the model output over the full 2D sphere to behavioural data. First, we describe the dynamic sound localisation model. Then we describe the localisation experiment and compare the results obtained here to the model data. Finally, we adjust a subset of the parameters to investigate their effect on localisation performance. The current experiments focus on small head rotations (10°) along the yaw axis. Small head movements have been shown to comprise the majority of natural head movements [17], and can be considered, if necessary, as a first step in a more complex movement framework. An added benefit of small rotation angles is that they allow us to approximate the rotation with a single axis and a constant acceleration. All experimental data, including the localisation model, are publicly available in the Auditory Modeling Toolbox (AMT) [18].

## Methods

### Template definition

The proposed model utilises a ‘template-matching’ procedure which requires an acoustic template **T**_*A*_ that the observed information is compared to. We assumed that **T**_*A*_ is the acoustic ‘knowledge’ that the brain has learned and stored over a lifetime of experience, and is thus signal-independent.

To compute **T**_*A*_, a set of acoustic features was extracted from the subject’s head-related transfer functions (HRTFs) and the signals received at each ear. This process is identical to the feature extraction described in earlier work [19].

The ITD template *T*_*itd*_ was computed as the difference between times of arrival (TOAs) of the head-related impulse responses (HRIRs) at each ear. The TOA was defined as the time it takes for the HRIR to reach a value 10 dB below its maxima. Each HRIR was low-pass filtered at a cutoff frequency of 3000 Hz before deriving the TOA. Then, the ITDs (in time units) were transformed into a scale of just-noticeable difference (JND) units, such that the error on the ITD was modeled as an additive instead of multiplicative factor (for further explanation, see [19]).

Next, we consider the spectral cues for the left **T**_*L*_ and right **T**_*R*_ ear separately. For each, the HRIRs and the stimulus were passed through a Gammatone filterbank with 32 channels spread across the frequency range of 300 Hz to 15 kHz in equivalent rectangular bandwidths (ERBs) following. These processed signals were half-wave rectified, low-pass filtered using five sequential first-order infinite impulse response (IIR) filters with a cut-off frequency of 2000 Hz, and then transformed to a logarithmic domain (in dB). This stage simulates a simplified processing of the inner hair cell [20]. The filterbank outputs were then summed over time per frequency channel. Thus, **T**_*L*_ and **T**_*R*_ denote vectors with monaural spectral information in dB along the ERB channels.

Ultimately, the ITD and the monaural spectral vectors for both ears are combined into **T**_*A*_ which is a matrix containing the combined vectors per template source direction:

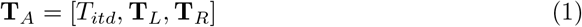

In this article, **T**_*A*_ consists of 2042 directions that are uniformly distributed over the sphere. These directions were obtained through spherical-harmonics interpolation of the measured HRTFs, involving Tikhonov regularisation to account for measured directions not covering the full sphere [21].

### Generative model

We assume that the listener wants to determine the source direction based on all prior information about the environment and all sensory information collected during the head movement. This prior and sensory information is combined into a posterior probability density function (PDF), from which finally a point estimate is retrieved.

To explain this process step-by-step, we first introduce the function of the spatial prior *p*(*ψ*). Then we discuss how the likelihoods of the acoustic information *L*_*A*_ and sensorimotor information *L*_*H*_ are computed. Finally we combine these factors into the posterior PDF and obtain an estimate of the sound source direction from this distribution.

### Spatial prior

In the Bayesian framework the probability of an occurring event may be affected by prior knowledge about the event. The spatial prior *p*(*ψ*) quantifies the listener’s a-priori assumptions about the source location before taking any sensory information into account. The vertical direction estimation appears to rely strongly on the involvement of a spatial prior, whereas azimuth estimation does not seem to be influenced by a prior much [22]. This can be modelled with a Gaussian spatial prior around the horizontal plane with a limited SD of about 12°. However, the best fitting SD of the prior seems to depend on the decision rule used.

The spatial prior is only one example of possible prior information available to a listener. Priors can be related to any variable, such as the number of sources [23], the movement properties of the sound source [24] or its spectral content [25]. In fact, the proposed model inherently relies on the assumption that the source spectrum is known and that the source is completely static in space.

### Acoustic sensor model

The acoustic sensor model compares the stored template information **T**_*A*_ to a vector of acoustic features present in the observed sound signal, **y**_*A*_, which consists of the noiseless ‘true’ state of the acoustic information, **X**_*A*_, corrupted with noise due to uncertainties within the auditory system or caused by the environment:

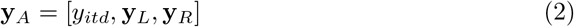

with

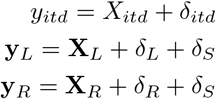

where *δ*_*itd*_ is the error on the ITD measurement with variance *σ*_*itd*_, *δ*_*L*_ and *δ*_*R*_ are the errors on the left and right monaural spectra measurements, respectively, and *δ*_*S*_ is the error due to imperfect knowledge of the sound source (assuming a central process, this is the same at both ears).

With that, we define the full covariance matrix of the acoustic cues as:

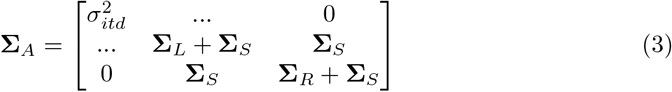

with

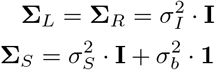

where the size of the identity matrix **I** is given by the number of frequency channels, 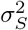 is the expected channel-dependent variance on the source, and 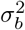 is the expected variance on the overall intensity of the source. All error terms are assumed to be zero-mean Gaussian noise. Note that we assumed the spectrum to be time-invariant and the source in the far field.

We then consider the acoustic sensor model:

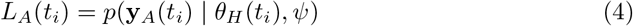

where **y**_*A*_(*t*_*i*_) and *θ*_*H*_ (*t*_*i*_) are the observed acoustic information and the true head orientation, respectively, at time-step *t*_*i*_, and *ψ* is the true sound source direction, which here is assumed to be static.

The expression in Eq 4 is calculated by computing the Mahalanobis distance between the measured acoustic cues **y**_**A**_(*t*_*i*_) and the set of acoustic cue templates **T**_**A**_ and covariance matrixΣ_*A*_. This is done at each sampled sound source direction, given the current head orientation *θ*_*H*_ (*t*_*i*_).

### Motor sensor model

The motor sensor model is defined as:

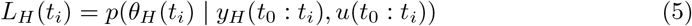

where *θ*_*H*_ (*t*_*i*_) and *y*_*H*_ (*t*_*i*_) are the true and observed head orientations at each time step *t*_*i*_, respectively, and *u* is the motor command signal, which is represented by the speed *ω*(*t*_*i*_) of rotating the head around a given axis. These variables are defined as:

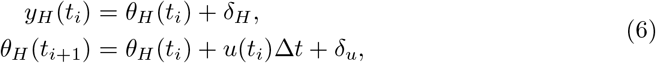

The additive noise on both the movement equation and the sensor equation is again assumed to be zero-mean white Gaussian noise *δ*_*u*_ *∼ 𝒩*(0, *σ*_*u*_) and *δ*_*H*_ *∼ 𝒩*(0, *σ*_*H*_). Thus, *σ*_*H*_ describes the noise on the head orientation observation at each time step and *σ*_*u*_ describes the noise on the motor command that steers the head.

Assuming head orientation measurements to be independent of acoustic measurements, we show in [16] that Eq 5 can be reformulated. The dependency on all sensor readings and all head rotations executed so far can be expressed recursively as

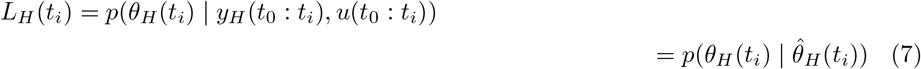

with 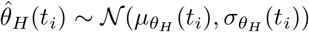 the estimated head orientation updated at each step through a Kalman filter with:

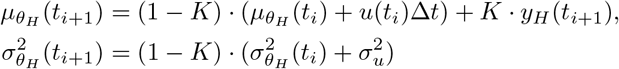

and K the Kalman gain:

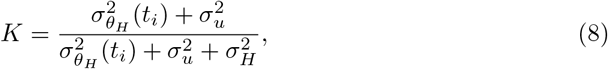

The expression in Eq 7 is calculated by computing the Mahalanobis distance between given head orientation *θ*_*H*_ (*t*_*i*_) and 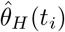 .

At the initial time step *t*_0_ we define: 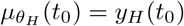 and 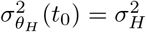.

### Posterior computation

By marginalisation over all possible head orientations and using Bayes’ theorem, we can combine the spatial prior and sensor model output to obtain the joint posterior PDF:

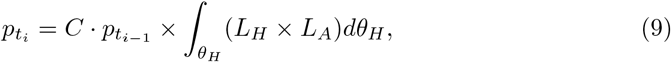

Turning to Bayesian terminology, 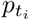 is the posterior PDF, 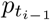 is the prior PDF and the joint sensor model computes the likelihood. *C* is a normalisation constant. Note that the prior at time step *t*_*i*_ equals the posterior from time step *t*_*i−*1_. At the initiation of the recursive process, 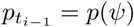, which is the spatial prior. The detailed derivation of this equation is explained in [16].

### Localisation decision rule

Eq 9 returns a probability distribution over the sphere, i.e., a probability from a large but discrete set of source directions. The last step in the process is to obtain a point estimate from this posterior PDF. To do so, a decision rule must be defined. The present model uses the posterior matching (PM) strategy, where a weighted random sample is taken from the posterior PDF. However, it is easy to implement other strategies, such as the maximum a posteriori (MAP) strategy, i.e., selecting the location at the maximum of the posterior. It was found that localisation performance lies somewhere between the MAP and PM strategies [22].

## Model parameters

The model noise parameters were set based on generalised (not listener-specific) findings from previous behavioral experiments. Table 1 shows the generalised parameters used in our study, of which the derivation is described next.

**Table 1.** Noise parameters and their chosen values.

| Symbol | Noise imposed on: | Value |
| --- | --- | --- |
| $\sigma_{itd}$ | ITD measurement | 0.6 JND |
| $\sigma_I$ | Spectrum measurement | 3.5 dB |
| $\sigma_S$ | Source spectrum knowledge | 0.0 dB |
| $\sigma_b$ | Source amplitude knowledge | 10.0 dB |
| $\sigma_p$ | Spatial elevation prior | 29.6° |
| $\sigma_H$ | Head orientation measurement | 0.0° |
| $\sigma_u$ | Head motor command | 0.0° |

### General

The stimulus was a broadband time-invariant Gaussian white noise burst. The duration was varied between 5 and 1000 ms, in order to test the effect of duration.

The template **T**_**A**_ was listener-specific, i.e., derived from the individual’s measured HRTFs. Earlier work suggests that the auditory system processes information (ITD [26], ILD [26, 27], spectrum [25, 28]) on a short time scale of about 5ms. For this reason, the time step size for the recursive updating of the posterior was set to 5ms.

The head movement was simulated as a yaw rotation with size *α* = 10°. The head movement trajectory followed a quadratic curve with constant acceleration 20°*/s*^2^, similar to the average trajectory of the experimentally measured head movements.

The spatial prior was assumed to emphasise directions around the horizon [22, 29]. That is, for elevation, the prior has a mean around zero and a restricted variance *σ*_*p*_. In [22], the optimal value of *σ*_*p*_ was found to be around 11.5°. However, this value was determined from a localisation experiment that only included source directions in the elevation range of [*−* 35°, 35°] in the frontal hemisphere. To extend the range to elevations up to 90°, *σ*_*p*_ was scaled up to 29.6°.

The localisation task was simulated for 33 source directions and repeated 20 times without head movement, 20 times with a 10° rotation to the left, and 20 times to the right. This reflects the sound directions and repetitions used in the behavioral experiment. The simulations were repeated for each subject that participated in the behavioral experiment, then the results were pooled for further statistical analyses.

### Acoustic information

The SD on the ITD measurement at each time step *t, σ*_*itd*_, was set to 0.6 JND. These values and units were derived earlier for the static localisation model [19]. In the discussion the effects of higher ITD measurement noise are reported.

We calculated the SD on the measurement of the spectral content at each time step *t, σ*_*I*_, with the same equation that was used for the static localisation model [19]:

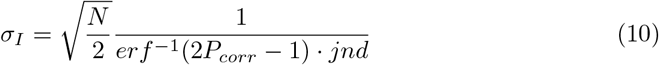

where *N* is the number of frequency channels (in this case 32),*P*_*corr*_ is the probability of correct responses and *jnd* is the just-noticeable difference measured.

Because this model computes the posterior recursively, as opposed to a single computation, and spectral information is found to be integrated over time [30, 31], *σ*_*I*_ needed to be adjusted to compensate for the number of looks made during stimulus presentation.

For a stimulus of 500ms, the JND corresponding with a 79% probability of correct responses was found to be 0.5 dB [32]. Inserting this in Eq 10 returns *σ*_*I*_ = 3.5 dB. Assuming that each time step contains the same noise and one look is made every 5ms, *σ*_*I*_ needs to be multiplied by 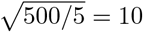 to obtain a posterior equivalent to one from a single look.

We tested this using the same conversions and findings for an experiment conducted with a 250ms stimulus (*jnd* = 0.65 dB,*P*_*corr*_ = 75% [27]). We then obtained from Eq 10 *σ*_*I*_ = 5.5 dB, which is indeed relatively close to 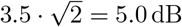.

As the sound stimulus was a broadband noise with no variation in its spectral content between trials and the subjects went through an initial training of 300 trials, we assumed that the subject had precise knowledge of the stimulus spectral content.

Therefore, the SD on the knowledge of the spectral content of the incoming sound source at each time step *t, σ*_*S*_, was set to 0.0 dB.

### Sensorimotor information

The SD on the measurement of the head orientation at each time step *t, σ*_*H*_, and the SD on the motor command steering the head rotation, *σ*_*u*_, were both initially set to 0°. In other words, this assumed that the listener can perfectly estimate and control the head orientation. Human subjects are able to report motion and orientation perception with very high precision [33, 34]. In a seated position, the standard deviation of head rotation around the starting position (notated in our model as 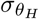) is around 2° [11]. In the discussion the effects of higher motor noise are reported.

As with virtually all sensory systems, motor imprecision increases with stimulus magnitude [35, 36], i.e., noise increases with exerted force. However, the acceleration is kept constant in the model rotations, so motor noise (*σ*_*H*_ and *σ*_*u*_) can be assumed here to be additive.

### Acoustic measurements and behavioural experiment

#### Apparatus

The behavioural localisation experiment and acoustic HRTF measurements were conducted in a semi-anechoic room with 91 speakers (E301, KEF Inc.) distributed over the sphere within the elevation angles *−*47° to 90°. A head-mounted display (HMD, Oculus Rift, CV1, Meta Inc.) was used for the visual presentation of virtual reality (VR) and three infrared cameras were used for the tracking of the listener within the six degrees of freedom.

The experiment was controlled by a computer running a 64-bit Windows 10, equipped with an 8-core, 3.6-GHz CPU (i7-11700KF, Intel Inc.), 16 GB of RAM, and a graphic card with dedicated 8 GB of RAM (GeForce RTX 3070, NVIDIA Inc.). The experiment was controlled by the ExpSuite 1.1 application LocaDyn, version 0.9.7.

The tracking system provides a translation accuracy of below 1 *cm* [37] and a rotation accuracy of below 1° (for a similar tracking system, [38]). The position and orientation of the subject’s head were recorded for later analyses.

#### Subjects

Eight normal-hearing subjects (four female, four male) participated in the experiment. Their absolute hearing thresholds were within the average (*±*1 standard deviation, SD) of the age-relevant norms [39, 40] within the frequency range from 0.125 to 12.5 kHz. The age range of the subjects was between 22 and 33 years.

#### Stimuli

The acoustic stimulus used in this experiment was a wideband (20 to 20000 Hz) white noise burst, gated with a 5-ms cosine ramp. Each trial used the same noise realisation, thus there was no variation of the spectral content between trials. The duration was 500 ms for the static condition and was gated off after 10° of head rotation for the dynamic condition (811*±*241 ms). Presentation level was measured to be 48 dBA SPL at the ear drum, with a *±*5 dB level roving range between trials.

Stimuli were played over loudspeakers using vector base amplitude panning (VBAP) [41]. Thirty-three source directions were distributed over the full sphere, which are visible in Fig 1 and Fig 2.

**Fig 1.**
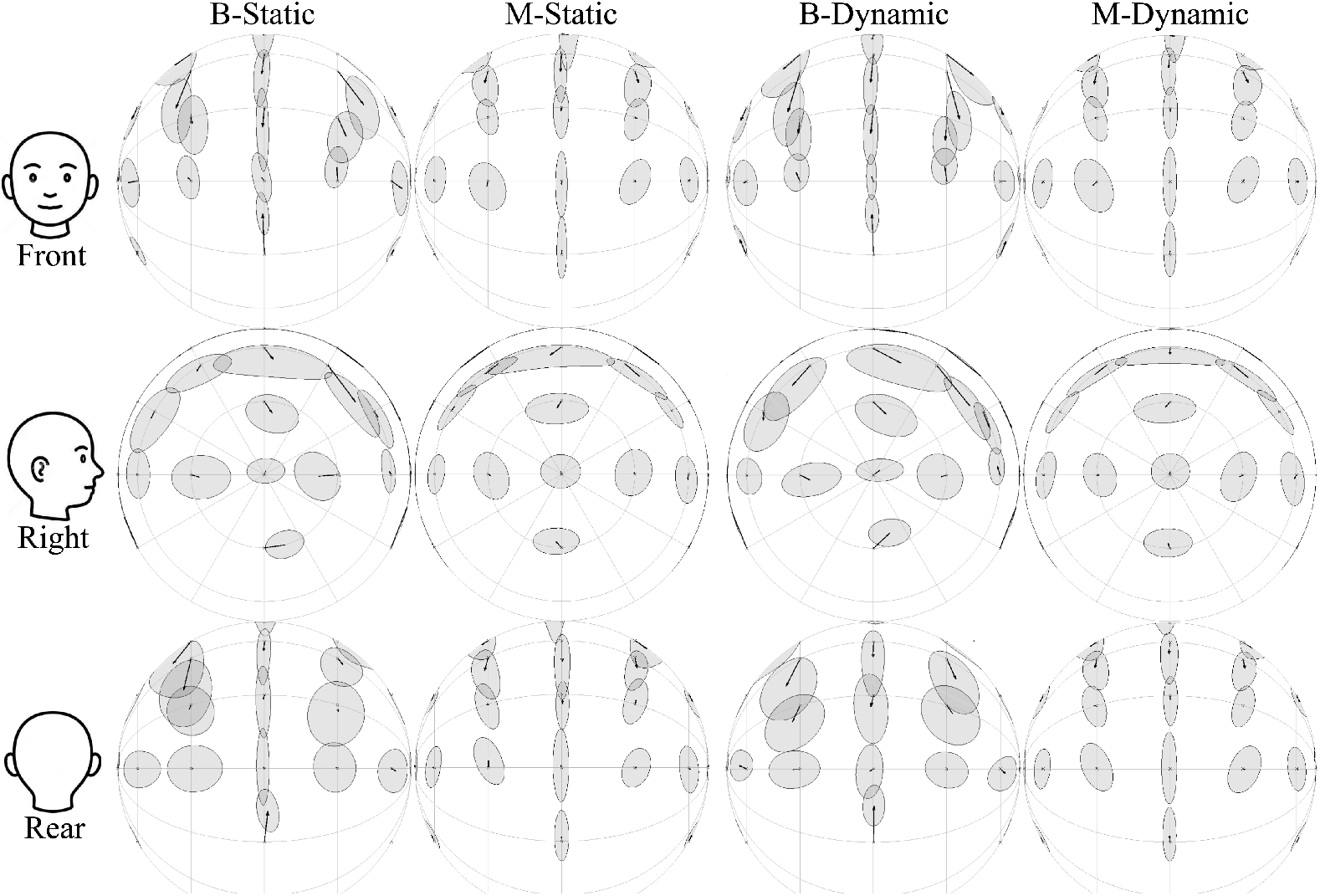
Centroids and Kent distributions of behavioural and modelled responses in the static and dynamic conditions, averaged over eight subjects. The rows show the same data viewed towards the front, the right, and the back of the head. Front-back confusions are excluded.

**Fig 2.**
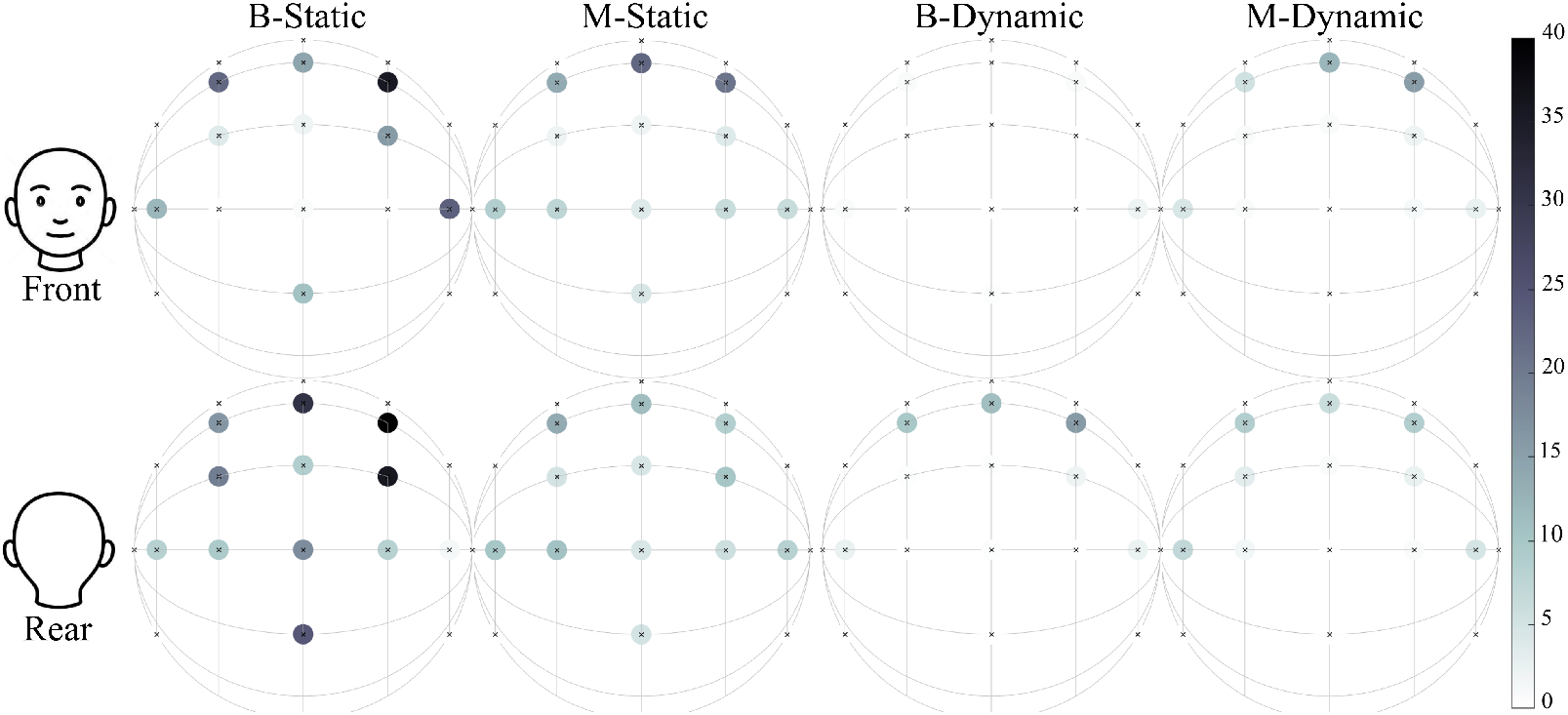
Front-back confusion rates [%] of behavioural and modelled responses in the static and dynamic conditions, averaged over eight subjects. The rows show the same data viewed towards the front and the back of the head.

#### Procedure

The localisation task procedure was identical to that of [7]. At the start of each trial, the subject kept their head still on the reference orientation at (0°, 0°). The stimulus was then played and the subject remained still or initiated rotation depending on the movement condition. At the end of the stimulus, the subject pointed towards their perceived source direction with a hand-tracking device to provide their localisation estimate. No feedback was provided about performance during or after the trials.

In the condition labelled ‘static’, the subject was instructed to keep the head still for the duration of the stimulus. Trials which exceeded 2° of movement in any direction were excluded. For the condition labelled ‘dynamic’, the subject was instructed to make a single-sided rotation (either to the left or to the right) as soon as they heard the stimulus onset. Half of the trials instructed a leftward rotation, the other half was rightward. The head rotation speed was unrestricted, but was monitored through the tracking system of the VR headset and recorded for analysis.

In total, the static experiment consisted of 660 trials (33 directions and 20 repetitions) per subject. The dynamic experiment consisted of 1320 trials per subject: 660 for a leftward rotation and 660 for a rightward rotation. The trials were divided into 6 blocks, with trials and blocks presented in random order.

Subjects were trained before commencing the experiment. The training consisted of 300 trials with the 500ms white noise burst spatialised from a direction randomly selected from a uniform distribution over the available directions of the subject’s individually measured HRTF set. Subjects were not excluded based on their performance at the end of training.

### Localisation metrics

There are many metrics available for sound localisation performance, this makes comparison between localisation studies difficult. It is, however, generally accepted that a distinction needs to be made between two types of errors [15]. The first type is the local error. Here we use the lateral-polar coordinate system [42], (*θ, ϕ*), where *θ∈* [*−*90, 90] and *ϕ∈* (*−*180, 180], with (*θ, ϕ*) = (0, 0) defined as straight ahead. The local error was expressed in root mean-squared error (RMSE) value of the lateral and polar errors. Polar errors larger than 90° in magnitude were excluded from the local errors [43].

The second type of error is the reversal error, which generally is reported as a percentage, i.e., the rate of reversals in a given set of trials. The first reversal error considered was the quadrant error (QE) rate, which is defined as any polar error larger than 90°. Additionally, we used the front-back confusion (FBC) rate and the up-down confusion (UDC) rate, which are defined as any response crossing the frontal plane and the horizontal plane, respectively. This is the same definition as used by Carlile et al. [15], and thus allows for a direct comparison between present and previous data. Note that this is a very coarse definition for the reversal error, as it confounds FBCs and UDCs with local errors near the frontal or horizontal plane, respectively.

## Results and discussion

### Global statistics

First, we discuss the global results, in order to compare to the existing literature. Next, a spatial analysis is presented in order to quantify directional dependence. Finally, a number of significant model parameters were varied to test their effect on predicted localisation performance.

### Behavioural data

Table 2 presents the localisation data of both the behavioural experiment and the model simulations (means and SDs averaged over the subjects), for static and dynamic conditions. For the static condition, the means and SDs of the behavioural results were similar to those from [43, 44]. The slightly lower SDs might be an effect of our smaller number of subjects and higher number of responses collected per direction.

**Table 2.** Averages and SDs of behavioural and modelled localisation performance in the static and dynamic conditions. The performance is represented as the lateral and polar RMSE (in degrees), QE, FBC, and UDC rates (in %). Means and SDs were computed over eight (virtual) subjects. For comparison, the stimulus duration is reported along the results from [43, 44]. N.R.: not reported.

| Condition | Dur.<br>(ms) | L. RMSE<br>(deg) | P. RMSE<br>(deg) | QE<br>(%) | FBC<br>(%) | UDC<br>(%) |
| --- | --- | --- | --- | --- | --- | --- |
| B-Static [43] | 300 | 10.6 ± 2.0 | 22.7 ± 5.1 | 4.6 ± 5.9 | N.R. | N.R. |
| B-Static [44] | 300 | 12.4 ± 2.2 | 21.9 ± 2.4 | 17.1 ± 5.6 | 12.8 | N.R. |
| B-Static | 500 | 8.1 ± 1.4 | 25.1 ± 3.2 | 7.9 ± 4.5 | 10.7 ± 6.3 | 6.7 ± 4.2 |
| B-Dynamic | 500 | 8.1 ± 1.7 | 20.5 ± 5.2 | 0.7 ± 1.1 | 1.6 ± 1.4 | 4.2 ± 4.1 |
| M-Static | 100 | 4.0 ± 0.3 | 24.0 ± 1.2 | 4.7 ± 2.1 | 6.0 ± 1.8 | 2.6 ± 0.9 |
| M-Dynamic | 100 | 4.1 ± 0.1 | 20.8 ± 1.7 | 1.6 ± 0.6 | 2.7 ± 0.9 | 2.2 ± 0.7 |

In the dynamic condition, the FBC rates improved among all subjects, regardless of localisation performance found within the static condition. This is similar to [7] and an indication for the significance of the dynamic cues obtained from yaw rotation.

Yaw rotation also decreased the UDC rate. This suggests that spectral cues above and below the listener vary differently with yaw angle. However, it is also possible that this reduction was caused by a roll-component in the yaw rotations made, which helps distinguish between the lower and upper hemispheres [16]. Indeed, subject 4, who showed the largest decrease in UDC score with movement, performed roll rotations as large as 8° during instructed yaw movement.

Finally, there was also a decrease in polar error in the dynamic condition. This can be attributed to the calculation of the polar RMSE, which still allows some localisation estimates in the opposite hemisphere to be included [43]. When omitting all estimates in the incorrect hemisphere (i.e., omitting FBCs instead of QEs) the decrease in polar error disappeared. This also agrees with previous findings [7].

### Model data

The model localisation errors are reported for 100 ms. For this duration, the mean errors were similar to that of the behavioural data. However, the model over-performed for 500 ms stimuli. The effect of the stimulus duration on localisation performance is further discussed below.

With focus on the 100 ms simulations, the polar errors and quadrant errors have similar values to those of the behavioural data. The dynamic condition also removed most QEs and FBCs, as was seen in the behavioural data. The FBC rate in the static condition was lower for the model. However, the high standard deviation in the behavioural reversal errors shows that this phenomenon is highly subject dependent. The lateral errors were quite small. This may have been the result of *σ*_*itd*_ being set too small. The effect of the ITD noise on the model performance is tested below.

Finally, the SDs between model ‘virtual listeners’ are small compared to the behavioural results. This is not surprising, as the individual differences between subjects are likely not fully explained by the individual HRTFs. The same noise parameters were used for each individual, while it is likely that they differ per individual [45]. Furthermore, higher level processes such as listening strategies [46] or attention [47] will also be a cause for individual differences.

## Spatial analysis

Fig 1 visualises the behavioural and model localisation results per source direction on the sphere around the listener. Following the methods of visualisation of previous work, the localisation responses were modelled as elliptical Kent distributions [15, 48].

The centroids visualise the bias, i.e., the mean vector, of all responses for one source direction. The ellipsoid outlines visualise the equal probability contours of the distribution of responses. The major and minor axes of the ellipsoid are two SDs in length and represent the first two orthogonal ‘principal components’ of the dataset that account for the maximum amount of variance in the data. FBCs and UDCs were excluded from the local error analysis, FBCs were visualised separately in Fig 2.

### Kent distributions

The direction of the centroids of the behavioural results in Fig 1 suggest a spatial prior centred around the horizontal plane; this is consistent with earlier findings [22]. There also is a slight bias away from the median plane, towards the left and right ear. Several studies have shown that humans during sound localisation display a peripheral bias in their responses that increases with eccentricity [49–51], although this is not the case in all studies [22]. The data shows a third bias for sources on the frontal plane (polar angle *±* 90), towards the frontal hemisphere. This may be related to the distribution of reversal errors, which shows more back-to-front (B2F) than front-to-back (F2B) errors. This will be further discussed in the next section. Two explanations for spatial biases have been proposed previously: 1) a Bayesian approach describes a pre-existing spatial prior for certain locations that is added to the sensory information, 2) alternatively, responses may be pulled to certain directions due to compressions and expansions in the sensory representations of auditory (and visual) space [51].

Fig 1 shows that the selected spatial prior for the model results in vertical biases that are similar to the behavioural data, this is evidence that the spatial biases can, at least in part, be explained by a prior. Due to the Gaussian shape of the spatial prior, sources at higher elevations are affected more heavily than those around the horizon. It is possible that a different prior shape may better capture the behavioural data, this is investigated below.

The biases did not change in the dynamic simulations, whereas the biases increased in the behavioural results. This suggests that rotation increased the uncertainty on the acoustic information, which increased the reliance on the spatial prior. Possibly, this effect was not be seen in the model because the head orientation was perfectly known. Below we investigate the influence of increased sensorimotor noise.

The model Kent distributions in Fig. 1 are also very similar to those of the behavioural data. This means that, for most source directions, the spread in responses can be predicted. The most frontal source directions on the horizon are an exception, as the spread here was higher than in the behavioural results. A stronger spatial prior might change this.

The spread in model responses for sources behind the listener were nearly identical to those in front. On the contrary, the behavioural data contained a much larger spread to the rear. This suggests that the increase in variance, i.e., decrease in precision, found in the rear hemisphere of the behavioural results cannot be (fully) attributed to a lower spatial resolution in acoustic cues. Instead, a response or ‘pointing’ error may have been responsible. Even if the auditory system can perfectly estimate a source location, the action of pointing in a direction as a response may introduce an additional error. It is reasonable to hypothesise that sources behind the listener are more difficult to point towards consistently. Pointing errors have been modelled previously [45, 52], but more research is required to accurately model the spatial dependence of a response error.

Comparing the movement conditions, the model spread decreased with head movement for sources above the listener. This change is not visible in the behavioural results, and can be explained by the increased spatial information obtained from dynamic cues which overrule the spatial prior. This is evidence that perfect knowledge of the head orientation in the model made the acoustic cues more informative than in the behavioural experiments.

The centroids and Kent distributions are generally symmetrical around the median plane for both movement conditions in both the behavioural and model data. We defined a metric for symmetry of the bias vectors as the Euclidean distance *D* between the vectors on the left hemisphere [*x, y, z*]^*L*^ and the vectors on the right hemisphere, mirrored over the median plane [*x, −y, z*]^*R*^. This metric revealed that the behavioural results for dynamic localisation (*D* = 0.055) are more symmetrical than the static condition (*D* = 0.106). The asymmetry in the static condition may be related to the asymmetry in FBCs, which were removed from the data. This will be discussed in the following section. The model responses were more symmetrical in both movement conditions (static: *D* = 0.026, dynamic: *D* = 0.022).

### Reversal errors

The behavioural data in Fig 2 reveal that B2F confusions were more common than F2B confusions: 67.6% and 32.4%, respectively. Most notable are the source directions directly in front of the listener, which showed nearly no FBCs at all. The model FBCs were symmetrical: 51.1% and 48.9%. This suggests that non-acoustic factors influence the preferred hemisphere, e.g. a spatial prior towards the front. A frontal prior would also explain the bias vectors in Fig. 1, which point from the frontal plane towards the front.

However, previous studies have shown that the quantity and spatial distribution of reversal errors were highly subject-dependent. For example, Makous and Middlebrooks found two subjects with a significant majority of B2F confusions, two with a majority of F2B confusions, and two with no significant preference [14]. Similarly, this study contained three subjects with a B2F majority, one with a F2B majority, and four with no clear preference.

For static listening, there was also a left/right asymmetry in the FBC rates of the behavioural data. At high elevations, FBC rates between the left and right hemispheres differed by as much as 23%. Previous studies also found asymmetries in FBC rates between the left and right hemispheres [53, 54]. This asymmetry is, however, not found in the responses of all subjects [55, 56] and even may be gender-specific [57]. Many studies also found no significant asymmetry at all when averaging over all subjects [49, 56]. From the present findings and the available literature, it becomes apparent that reversal errors involve a complex process that differs between individuals.

### Effect of the stimulus duration

The model results revealed that stimulus duration was a significant factor in the localisation task. An inherent characteristic of recursive Bayesian inference is that, as long as new information is provided, errors will converge to zero. On the other hand, for humans, it has been shown that durations beyond 100 ms in static listening do not significantly improve performance [25].

Our auditory perception does not seem to process the immediately present acoustic information, but rather processes the outcome of temporal integration during a sliding window with a size of 150 to 200ms [30, 31]. In localisation studies along the horizontal plane, the azimuth estimation reached best performance for stimulus durations of only 3 ms [58]. In studies along the vertical plane, a longer duration of 80 ms was required to reach the best performance in the elevation estimation [25]. When head movements are allowed, a stimulus duration of approximately 100 ms seems to be required to provide a substantial benefit from the head movement [59].

Fig 3 shows that the model reached a performance plateau around 500 ms. This does not match the durations found in previous studies, and the plateau values are much lower too. This means that the current model does not yet correctly predict the temporal aspect of sound localisation.

**Fig 3.**
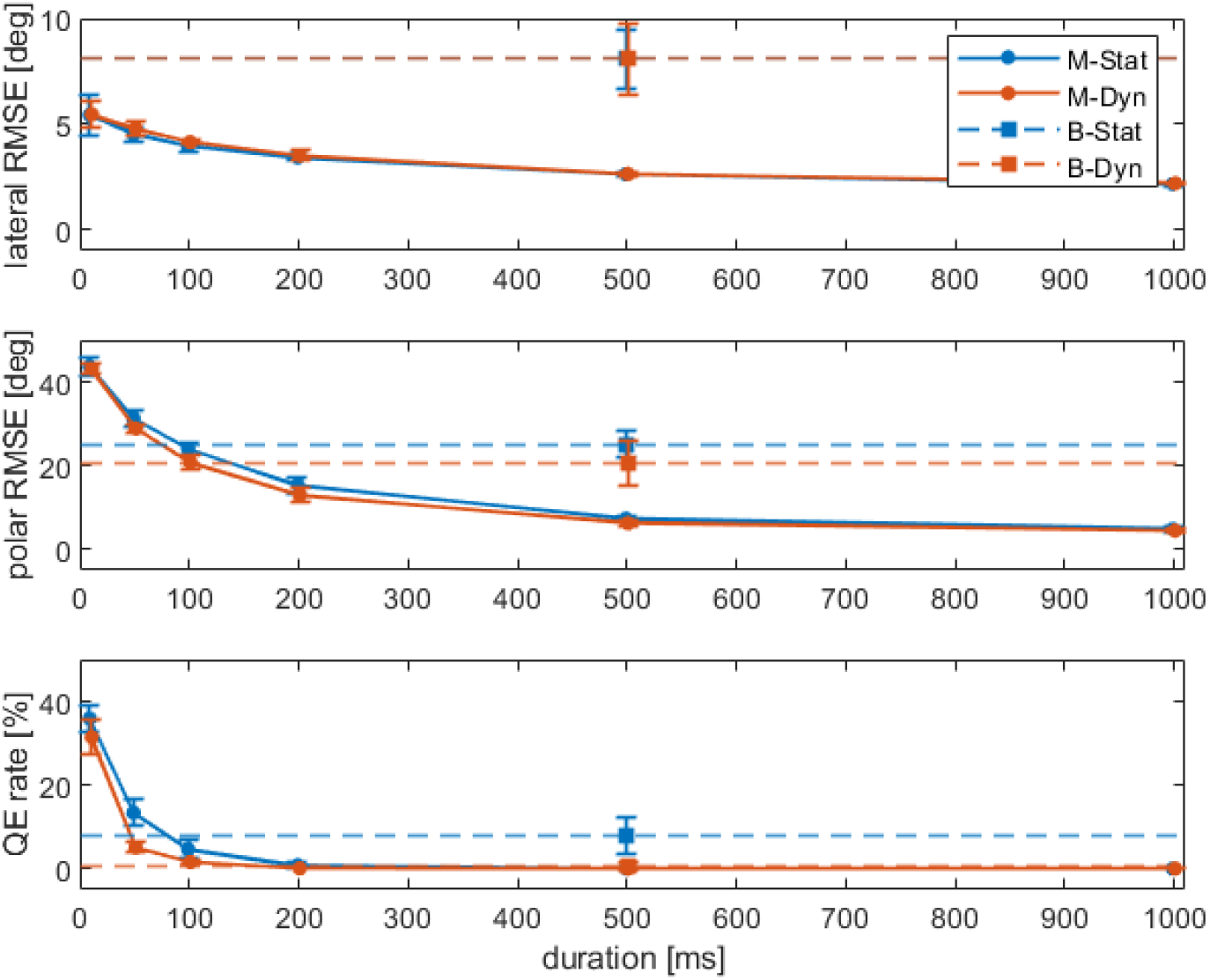
Lateral RMSE, polar RMSE, and QE rates of the modelled data as a function of stimulus duration. The symbols show the averages and the error bars represent *±*1 SDs over the (virtual) subjects. For reference, the horizontal dashed lines show the behavioural data obtained for the duration of 500 ms.

One solution could be to increase the update rate of 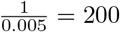 Hz to a higher rate, so that the plateau is reached quicker. There is evidence that the brain exhibits activity at such high rates [60], but it is somewhat unlikely that estimations about the world are updated at these same rates. Furthermore, this would only solve the duration taken to reach the plateau, not the low error values.

Alternatively, it may be that humans only optimally integrate acoustic information up to 100 ms of listening. Our auditory system might function as a leaky integrator, where not all acoustic information is stored for the estimation process [31]. The model data supports this, as the polar RMSE and QE rate cross the behavioural values between 100 and 200 ms. The use of a leaky integrator may be beneficial when we consider a dynamic environment. The present model assumes a single, static sound source. Assuming a possibly moving sound source would put a premium on forgetting old acoustic information as the state of the world would have changed in the meantime.

Interestingly, the lateral RMSE best matched the behavioural results at a single time step of 5 ms (5.5°). This suggests that the ITD information may not be integrated at all, and is instead remeasured and reevaluated entirely at each time step. Temporal changes need to be processed at such high rates that it may not be advantageous to integrate this information over time [61], especially when the sound source could be moving.

Earlier studies have determined two separate neural pathways for lateral and polar estimation [62, 63]. It is then possible that the pathway for lateral estimation works on a single look-by-look basis, whereas polar estimation integrates spectral information over time.

### Effect of the ITD noise

Table 2 shows that the lateral RMSE was too small compared to the behavioural results. Here, we tested new values of *σ*_*itd*_ to investigate whether this parameter is the cause of the discrepancy. The results were plotted in Fig 4.

**Fig 4.**
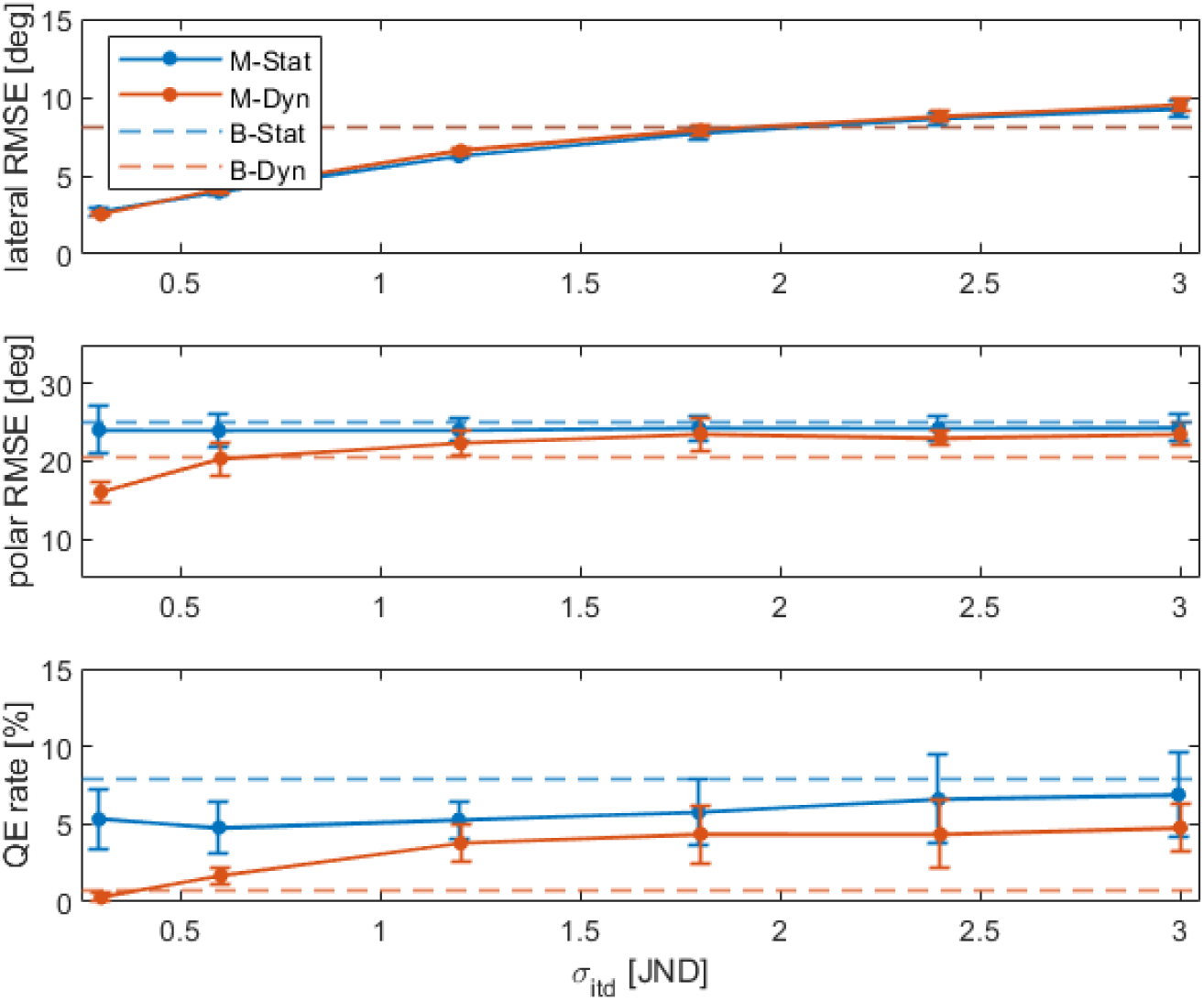
Lateral RMSE, polar RMSE and QE rate of the modelled data as a function of *σ*_*itd*_ (in units of the JND). The symbols show the averages and the error bars represent *±*1 SDs over the (virtual) subjects. For reference, the dashed lines show the behavioural averages.

Increasing *σ*_*itd*_ had a monotonic effect on the lateral error. Around *σ*_*itd*_ = 2 *JND* the lateral RMSE matched the behavioural data. However, this adjustment had a detrimental effect on the QE rates during dynamic sound localisation. As expected, polar errors remained mostly unaffected in the static condition, as the the ITD contains little to no information on the polar angle. Minimising *σ*_*itd*_ led to a complete removal of QEs in the dynamic condition. The reduction in polar RMSE, as stated earlier, was mostly related to the FBCs included in the calculation of polar RMSE. Together, this suggests that a low noise on the ITD cue is essential to utilise the dynamic cue that resolves reversal errors when moving the head. This also means that the ITD cue alone cannot account for the low lateral error.

### Effect of the sensorimotor noise

For the initial simulations, the head orientation was assumed to be perfectly known. Here we investigated the effects of an increased uncertainty on the head orientation. The simulations were rerun with different values for *σ*_*H*_ and *σ*_*u*_. The results were plotted in Fig. 5b. Fig. 5a illustrates the evolution of the head position information of the Kalman filter (Eq. 8) over 100 ms for *σ*_*H*_ = 5°.

**Fig 5.**
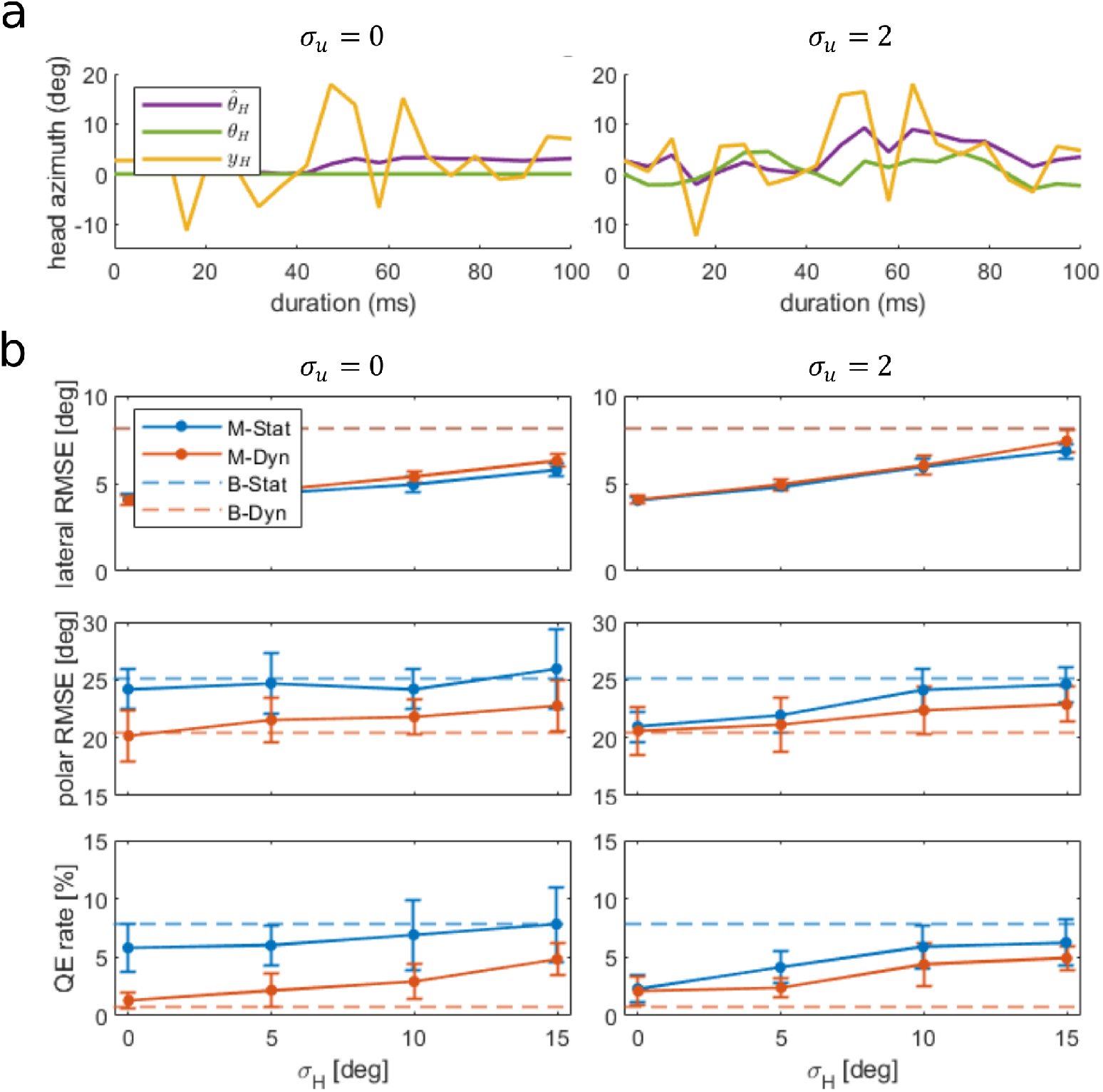
(a) Kalman filter input and output over 100 ms for *σ*_*H*_ = 5° and *σ*_*u*_ = 0° (left column) or *σ*_*u*_ = 2° (right column), where *y*_*H*_ is the observed head orientation, *θ*_*H*_ is the true head orientation and 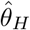 is the estimated head orientation, derived from the first two variables. (b) Lateral RMSE, polar RMSE and QE rate of the modelled data as a function head orientation measurement noise *σ*_*H*_ for head control noise *σ*_*u*_ = 0° (left column) and *σ*_*u*_ = 2° (right column). The symbols show the averages and the error bars represent *±*1 SDs over the (virtual) subjects. For reference, the dashed lines show the behavioural averages.

For *σ*_*u*_ = 0°, the filter was able to make a fairly accurate estimation of the head azimuth, even though the observations are noisy. For *σ*_*u*_ = 2°, the estimates were noisier, but so was the true head orientation. We see that for this small increase in variance in the motor noise the true head orientation *θ*_*H*_ started displacing up to 5° away from the initial position. This means that *σ*_*u*_ can realistically not be larger than 2° during static listening.

Interestingly, Fig. 5b shows that this movement improved localisation performance, as opposed to deteriorating it. It provided the model with sufficient acoustic information in the static condition that it performed similarly to the dynamic condition. This is evidence that incredibly small head movements can already provide the majority of dynamic cues to resolve reversal errors.

The plots show a monotonic effect between *σ*_*H*_ and the local errors and reversal errors. However, these effects are all surprisingly small, even for very high variances. Furthermore, these results suggest that the precision of human motor control is high, as the error metrics were most similar to the behavioural results at *σ*_*H*_ = 0° and *σ*_*u*_ = 0°. This means that it may be possible to make the assumption of perfect knowledge of the head orientation, significantly simplifying the dynamic localisation model.

Note that *σ*_*H*_ and *σ*_*u*_ will in reality be higher in the dynamic condition than in the static condition, as motor noise is multiplicative [35].

### Effect of the spatial prior

The bias vectors in the behavioural results appeared slightly stronger than for the simulations with *σ*_*p*_ = 29.6. This implies that a stronger spatial prior may be necessary. There are many different possible spatial prior shapes, e.g. prioritising the front or high lateral angles [64]. In this study the analysis was restricted to the horizontal prior. Fig. 6 shows the elevation gain [22] (i.e., the slope of a linear regression between responses and true directions) and QE rate for different values of *σ*_*p*_. As we are interested in the ‘pull’ towards the horizon, the elevation gain is a more appropriate indicator than the polar RMSE.

**Fig 6.**
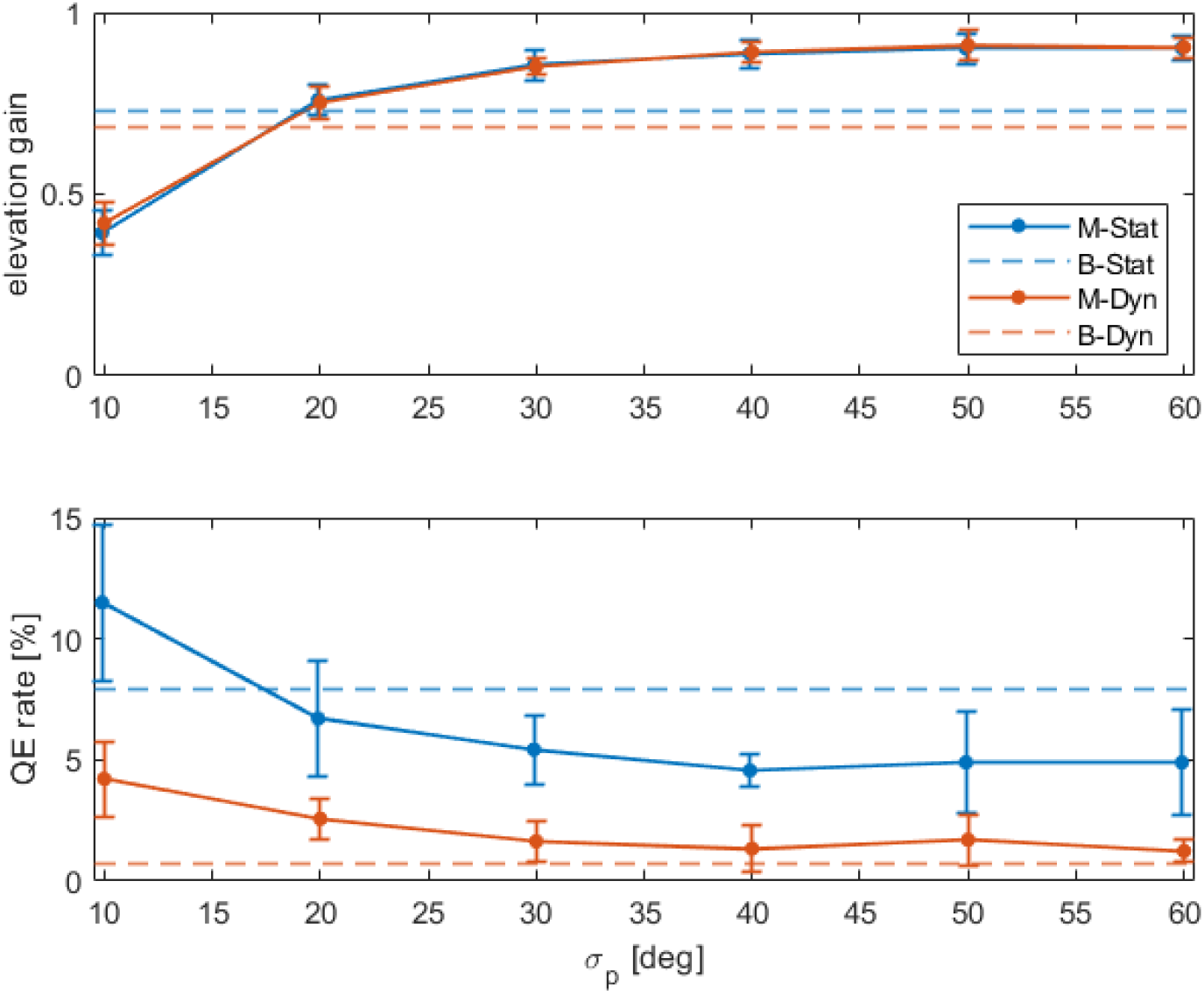
Elevation gain [22] and QE rate of the modelled data as a function of *σ*_*p*_ (in degrees). The symbols show the averages and the error bars represent *±*1 SDs over the (virtual) subjects. For reference, the dashed lines show the behavioural averages.

The plot suggests that the correct prior lies around *σ*_*p*_ = 20°, where the gain and the QE rates are closest to the behavioural results. As *σ*_*p*_ increases the prior approaches a uniform distribution and the responses approach a gain of 1. This shows that the bias present in human responses is not due to acoustic factors, but indeed due to a prior towards the horizon.

Next, it was noted that the prior affects sources at higher elevations more heavily than those around the horizon. There exist distribution shapes with sharper peaks and longer tails than the Gaussian distribution. One of such distributions is the Laplace distribution:

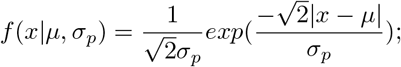

Fig. 7 shows how the response distributions change for the two different prior types, both with *σ*_*p*_ = 20°.

**Fig 7.**
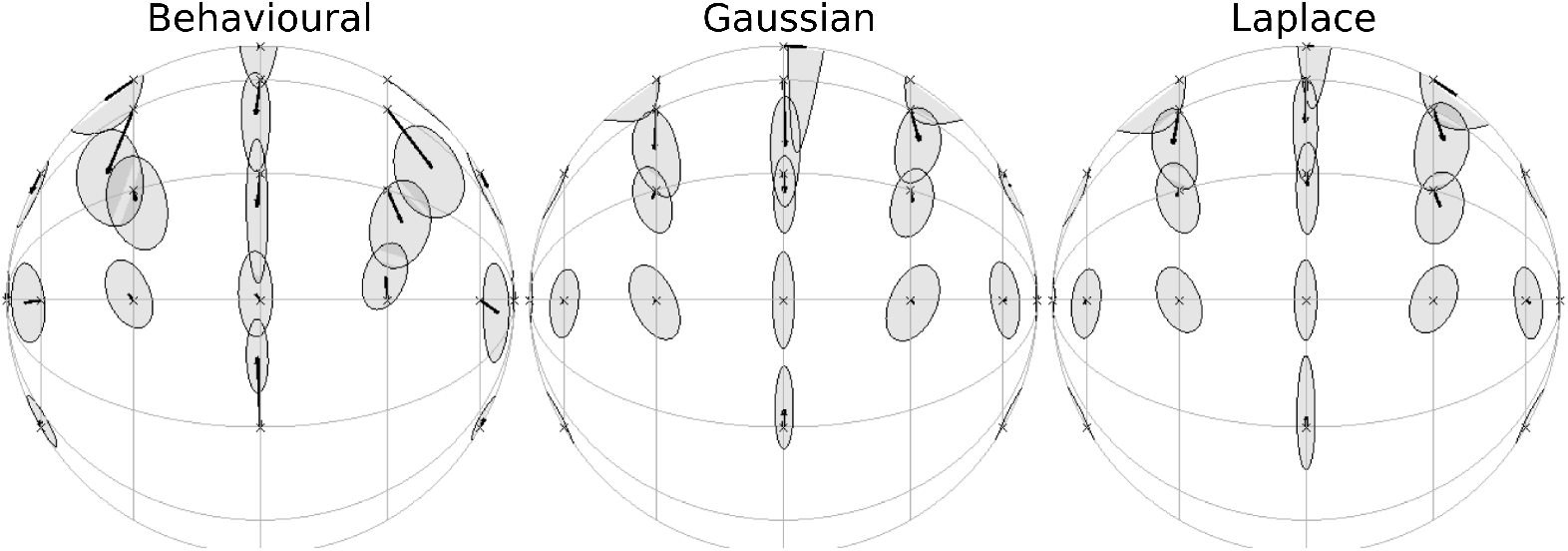
Centroids and Kent distributions of behavioural results, model output with a Gaussian prior and model output with a Laplacian prior (*σ*_*p*_ = 20°), viewed towards the front.

The Laplace prior seemed to work better than the Gaussian prior, especially on the median plane. The variances looked more similar to the behavioural results and the extreme spread on the source above the listener was removed. This is evidence that spatial priors are not necessarily Gaussian, but instead require a higher kurtosis.

## Conclusions

This article introduced a novel Bayesian model that allows for a bottom-up investigation of human performance during dynamic sound localisation. A localisation experiment was conducted in virtual reality to validate the model output.

With parameter values selected based on first principles, i.e., without the use of any fitting algorithm, it was shown that the model can predict and explain human performance quite accurately. This is an encouraging finding, which supports the hypothesis that model noise parameters can be derived from behavioural experiments. In a spatial analysis we showed in detail where the model agreed and deviated from the behavioural data. The largest differences were found for sources to the rear, which indicates that it may be necessary to model pointing errors in addition to the acoustic localisation task.

In addition, the effects of several model parameters were tested and reported. The duration analysis provided evidence against the assumption that humans can integrate acoustic information ad infinitum. Processing acoustic input as a leaky integrator may be beneficial when we consider a dynamic environment, as old positional information becomes less relevant as the state of the world changes. Adjustment of the ITD noise parameter revealed that the ITD cue alone cannot account for the lateral RMSE and QE rate found in behavioural data. Adjustment of the sensorimotor noise parameters suggested that humans can estimate and control the head orientation with high precision. Finally, adjustment of the spatial prior showed that this distribution may not be Gaussian, as a distribution with a sharper peak and longer tails better described the behavioural data.

The present analysis shows that this model can serve as a powerful framework to further investigate how the available cues and the way they are processed (e.g. with differing integration times or decision strategies) might affect human sound localisation, and ultimately help us better understand the complex system that shapes our perception.

## Acknowledgments

The authors would like to thank Philip Leong for providing the MATLAB code to the ‘Spak’ spherical data processing tool. Part of this code was rewritten for the purpose of this work and integrated into the AMT.

